# Controlling for false discoveries subsequently to large scale one-way ANOVA testing in proteomics: practical considerations

**DOI:** 10.1101/2022.08.29.505664

**Authors:** Thomas Burger

## Abstract

In discovery proteomics, as well as many other “omic” approaches, the possibility to test for the differential abundance of hundreds (or of thousands) of features simultaneously is appealing, despite requiring specific statistical safeguards, among which controlling for the False Discovery Rate (FDR) has become standard. Moreover, when more than two biological conditions or group treatments are considered, it has become customary to rely on the one-way Analysis of Variance (ANOVA) framework, where a first global differential abundance landscape provided by an omnibus test can be subsequently refined using various post-hoc tests. However, the interactions between the FDR control procedures and the post-hoc tests are complex, because both correspond to different types of multiple test corrections. This article surveys various ways to orchestrate them in a data processing workflow and discusses their pros and cons.

## 1. Introduction

In its most general form, the biomarker selection problem amounts to select some features that can be significantly associated to a difference of biological status, with the objective to subsequently refine decision making (diagnostic, prognostic, patient follow up, etc.). One of the most widespread and efficient ways to achieve biomarker selection is to perform a series of statistical tests (one for each feature available) and to retain those few that pass a user-defined significance threshold. This conceptual simplicity makes the procedure easily automatable, even though, on large omics data, two obstacles may raise questions and hinder the interpretation of the resulting biomarkers.

The first obstacle derives from the increase of instrumental throughputs: Many omics technologies provide quantitative measurements about thousands of features for each sample analyzed. From a biological perspective, this near exhaustiveness is a bliss, however, it also increases the chances that, for some of the putative biomarkers, random fluctuations concur with the change of biological status, hereby leading to artefactual significances; and thus to misleading biomarkers (a.k.a., false positives, a.k.a., false discoveries). To cope for this, it has become standard to control for the so-called False Discovery Rate (or FDR). FDR control procedures form a class of Multiple Test Corrections (MTCs), which aims at adjusting the significance value used to retain putative biomarkers as a function of the total number of measured features (see [1] for a proteomics oriented tutorial on FDR and related statistical notions). Therefore, in the end, the increment of omics measurement does not lead to an increment of false discoveries. Unfortunately, the theory underlying FDR control is not trivial and its correct application to biomarker selection remains at the center of an ever-renewed body of literature [2][3][4][5][6][7][8].

The second obstacle derives from the increasing complexity of the biological questions, and more specifically, from the increasing complexity of the associated experimental designs. In its simplest form, one seeks for biomarkers discriminating between two biological conditions, *e*.*g*., Healthy vs. Disease (in a clinical context), or Wild-type *vs*. Mutant (in a fundamental research context), e.g., [9][10]. However, more refined experimental designs are increasingly popular: Multiple comparisons (e.g., [9][11]), time-course (e.g., [12], which should not be confused with longitudinal analyses, for which specific temporal models are used [13]), strata (e.g., [14], where samples belong to patients with different degrees of illness, etc.), or multi-factorial designs (involving for instance a combination of severity, treatment and responsiveness, like in [12][15]). All these experimental designs share the need to compare *K* > 2 biological conditions. While an important corpus of applied statistic articles is dedicated to this subject, the most well-known statistical framework to address such experimental designs is that of the analysis of variance (or ANOVA). The reason for the popularity of ANOVA is twofold [16][17]: first, it is general enough to fit a variety of experimental designs; second, it generalizes the well-known t-test in an intuitive way making its interpretation easier. However, beyond this superficial simplicity, some technical difficulties appear, notably when Post-Hoc Tests (PHTs, see Section 2.3) are involved in the ANOVA.

If one depicts a dataset as a table where the rows correspond to the features (the putative biomarkers) and the columns correspond to the samples; then our first obstacle relates to the increment of the number of rows, while the second obstacle relates to the increasingly complex structure that binds the columns together. As previously mentioned, many solutions were proposed to each obstacle taken independently, among which FDR control (for the first obstacle) and ANOVA (for the second one) have been the preferred ones. Unfortunately, the combined application of FDR control and ANOVA is intricate as PHTs include a form of MTC that relates to the number of compared biological conditions. Broadly speaking, an FDR control amounts to a column-wise MTC, while a PHT amounts to a row-wise one. It is thus essential to orchestrate them in a manner that remains statistically rigorous and biologically relevant. This article reviews various ways to do so in a proteomics context (yet, its conclusions can easily be transposed to other omics settings).

## 2. Preliminary considerations

### 2.1. False discovery control in proteomic differential analyses

Because of the important inter-individual variance of the proteome, but also because of the classically small number of available assays, it is expected that at least a few proteins displaying differential abundance in the experiment do not convey any biological meaning (these proteins are the false positives of the biomarker selection process). Their precise proportion among the biomarkers selected is unfortunately not accessible. However, since Benjamini and Hochberg seminal work [18], we have known that this quantity can be estimated, using a False Discovery Rate (a.k.a. FDR, that is an abstract quantity that may differ from the real proportion of false discoveries in the data, but would be correct once averaged over multiple similar datasets [1]); or controlled at a given user-defined threshold (e.g., 1%), using adjusted p-values [19] (or similarly, q-values [20]).

Because of the complexity of the underlying theory, the user-defined control threshold is often conflated with the FDR itself, as in sentences like “Differentially abundant proteins were validated at 1%FDR”. In addition to advocate for a vocabulary simplification [1][5] with respect to biostatistics theory [21], this conflation tends to reduce the FDR framework to a feature selection method embedding its own multiple test correction. As such, the FDR control procedure has progressively stepped out of its original and prime role, namely assessing the statistical significance of a set of selected features (hereafter referred to as the “Statistical Rigor” role), and has gained two other roles.

The first additional role is that of a quality control measure: The tuning to a default value (typically 1% in label-free discovery analyses) is more justified by the standardization capacities it yields (hereby providing a ground level to compare different analyses from different labs) than by advanced statistical reasons. The second additional role is that of practical filter: It helps the researcher focusing on an acceptable number of proteins by filtering out the others.

Finally, FDR control is now motivated by three roles: Statistical Rigor (SR), Quality Control (QC) and Practical Filter (PF). Ideally, these three motivations concur: By filtering the list of putative biomarkers to 1%, hereby fitting with the QC standards, one brings rigorous statistical support to those deemed differentially abundant while substantially reducing their number to an order of magnitude that is compatible with the constraints of post-proteomic validation wet-lab experiments. However, every practitioner has ever faced a situation where this classical FDR threshold leads to a list of biomarkers, which is either far too short or far too long with respect to the anticipated validation experiments. Of course, it is possible to adjust the list by different tunings on preliminary filters (modifying the minimal number of peptides per protein, adjusting the minimal acceptable fold-change^1^, etc.), but if it is insufficient, one may be tempted to adjust the FDR threshold to unrealistically low or high thresholds. This illustrates well that concretely, the tuning of the cutoff often results from a negotiation between the QC and the PF roles, while the SR one is simply assumed inherent to the methodology. Although not absurd in a simple “binary” experimental design (*e*.*g*., Healthy vs. Disease) this view reaches its limit with more complex ones (time courses, strata, etc., see Section 1), which calls for a more explicit prioritization of the aforementioned roles.

### 2.2. ANOVA-related vocabulary

From a statistician viewpoint, the ANOVA is essentially a modeling framework, which assumes a response variable (e.g., protein abundance) to result (possibly after transformation) from a linear combination of explanatory variables (e.g., treatment, phenotype, experimental factor, etc.), up to some unexplained variations (a.k.a. noise, a.k.a. residuals). Under some mathematical assumptions, the noise can be decomposed in such manner that it informs us about the veracity of one of the hypotheses that prevailed when the practitioner built their linear model. More specifically, by comparing the noise decomposition to the F-distribution, one defines a statistical test, which can assess the significance of a contrast null hypothesis (a contrast null hypothesis is a hypothesis assessing a linear relationship between the response variables and the explanatory variables that equates to 0). For illustration purpose, let us consider the classical t-test, which null hypothesis reads:

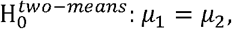

where *μ*_*i*_ is the mean of group *i*. Testing protein abundance *Y* as to know whether its means significantly differ between groups 1 and 2 is equivalent to considering the model,

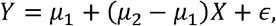

and then to test the contrast *μ*_1_ − *μ*_2_ = 0 When, *K* > 2 groups are considered simultaneously, a hardly more complex linear model makes it possible to test whether their *K* means equates, or not:

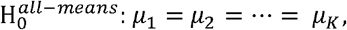

Such a test is referred to as an **omnibus test** and its null hypothesis can be recovered by testing simultaneously *K* − *1* well-chosen pairwise contrasts.

As most of the practitioners are seldom interested in the underlying mathematical model, it has become customary to conflate the modelling framework (the ANOVA) and the statistical testing of the contrast. For instance, the fixed-effect one-way ANOVA (i.e., an ANOVA model involving only one non-random experimental factor, as opposed to random- and mixed-effect models [25]) is often (and harmlessly) conflated with the omnibus test, and referred to as an “ANOVA test”. However, this conflation hinders the scrutiny of the interactions between the statistical tests and their multiplicity corrections, as it will appear in Section 3.2.

### 2.3. Post-Hoc Tests

The omnibus test is like any other statistical test: the interesting results it exhibits are those corresponding to the rejection of its null hypothesis (namely,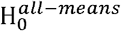), i.e., cases where the *K* groups can reasonably be suspected to have some of their means different from the others. For those cases, textbooks in statistics suggest to apply a post-hoc test (PHT) downstream of the rejected omnibus test, as to find which group means do not equate.

Essentially, a PHT is a tool that combines two distinct statistical steps. The first step is a series of other tests which aim at refining 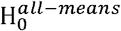. For instance, Tukey-Kramer HSD (Honest Significance of Difference) proposes to test *μ*_*i*_ = *μ*_*j*_, for all *i, j* ≤ *K* to find which pairs of groups display significant differences. Scheffé’s extends this by considering all possible subsets of groups. Conversely, Dunnett’s proposes to test only *μ*_*i*_ = *μ*_*j*_, for all 1 < *j* ≤ *K*, to check which groups are significantly different from the reference group 1 (see [26] for a thorough description of the available PHTs). The second step is a multiple test correction (MTC), to avoid that the inflation of *K* leads to an inflation of significant tests. For instance, if one compares a reference treatment to 100 alternative treatments that are in fact placebos, we can expect one of them to be significant by chance at a significance level of 1%. To cope for such effect, many PHTs encompass a Family-Wise Error Rate correction (FWER, [27]), that is a correction broadly akin to that of Bonferroni’s, despite subtle variations from one to another.

However, this workflow has two flaws that are usually not described in textbooks. First, significant rejection of 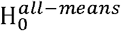 may not be confirmed by the downstream PHT and conversely, a significant rejection of a couple of means equality can be observed despite the non-rejection of 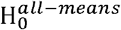 in the first place [28], which calls for alternative workflows [26]. Second, the conflation of the test series (those aiming at refining the omnibus test of 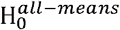) and the MTC into a single tool encourages a form of narrow-mindedness where statistical guidelines are blindly applied regardless of their relevance [29]. Let us illustrate this with two examples:

- What if one is only interested in the pairwise comparisons of the form *μ*_*i*_ = *μ*_*i*+1_ ? If one seeks proteins which are differentially abundant within all these pairs (without exception), one should not correct for multiple testing using an FWER.
- Conversely, if from the beginning, one is only interested in comparing the new treatments or placebos against the reference treatment, why should 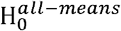 be tested in the first place? It would be equally sounded to consider as many pairwise t-tests as necessary, and then to apply a well-chosen MCT [30].

Naturally, the writers of most of the textbooks were aware of this, but we should keep in mind that their first editions have often been anterior to the big data era, or even to the desktop computer, as to propose guidelines to experimental research in various domains [29]. In those times, most of the computations were performed manually, or, at best, using a pocket calculator [31]. In such circumstances, proposing a straightforward workflow that minimized the computations (*e*.*g*., “spend time to deal with the pairwise comparisons only for the cases where an omnibus test has allowed you to reject 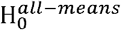) was probably more important than to account for the great diversity of experimental designs allowed by omics technologies (which were still to be created). Then, the weight of traditions did the rest [29] so that authors have only recently started to advocate for mindful use of PHTs [26][30].

### 2.4. Contrast-based testing at scale

In omics biology, Limma [32] is probably one of the most popular implementations of ANOVA. The reason essentially lies in its moderated statistics, which allows for a better estimation of the variance of the putative biomarkers in a context where biological samples may be scarce. This is why similar moderated statistics have been developed for count data (e.g., spectral counting, or any quantitative omics data produced by an NGS instrument), like EdgeR [33] or DEseq2 [34]. However, the influence of Limma goes beyond this.

Notably, it has popularized the test serialization, *i*.*e*., the testing of hundreds (or even thousands) of omics features (historically microarray probes) in an independent way. As opposed to the “pre-omics” statistical textbook guidelines, it does not systematically apply an omnibus test for each feature, and then a PHT in case of rejection. On the contrary, it allows for a direct focus on the specific contrasts of interest (as advocated in Section 2.3), together with the necessary MTCs.

Following this line has now become so mainstream that the warnings in Limma documentation ([35] Section 13.3) are classically ignored, while they pinpoint the difficulty of performing MTCs simultaneously on the rows and on the columns of the data table. To cope for them, the authors have proposed four options, all of which with restrictions on the condition of applications, or with an explicit lack of mathematical support (see Sections 3.2 and 3.3). However, these warnings step out of the specific case of Limma (and of its moderated statistics), as they are ubiquitous to complex experimental designs.

Finally, tools like Limma advocate for a rational separation between the various constituents of the one-way ANOVA workflow. Instead of considering a so-called “ANOVA test” (a linear model + an omnibus test) and PHT (including a row-wise MTC) among which FDR control has no specific identified place, one should think in terms of (*i*) a global linear model, (*ii*) the contrasts of interest, and (*iii*) the different types of MTCs needed. Owing to the difficulty of performing simultaneously row-wise and column-wise MTCs, this article proposes solutions to their application, in the specific case where the multiplicity of proteins tested is accounted for by an FDR; and the multiplicity of contrasts is accounted for by an FWER. Several options are discussed at the light of (1) the underlying biological motivation; (2) the type of linear model (classical or involving moderated statistics); and (3) the various roles classically endowed to FDR (statistical rigor, quality control and practical filter).

## 3. Various scenarios

This section is organized into four subsections, each presenting one family of scenarios. As most of them are rather classical (in the sense they are either derived from textbooks or from very popular statistical packages, e.g., Limma or multcomp [30]) the objective is essentially to discuss their pros and cons, depending on the proteomic question and on the three FDR roles (SR, QC and PF). Notably, we explain why the most classical scenario according to textbooks (Section 3.1) is of little interest in an omics biology context. Its natural alternative is depicted in Section 3.2. Because of an unquestionable statistical rigor, it can be over-conservative (i.e., it can discard too any putative discovery because excessively cautious), which may hinder the other roles (QC and PF). This is why Sections 3.3 and 3.4 describe various ways to loosen the rigor, while remaining sufficiently correct from a statistical viewpoint, at least according to the biological question and data analysis objectives. To illustrate the differences of conservativeness between these scenarios, they are compared on a publicly available dataset.

Before describing this dataset, some warning is necessary. Displaying a comparison of several statistical workflows on a given dataset is extremely slippery, for two reasons: First, it may give the wrong impression that any of the scenarios can be applied to any dataset. Although true from a computational standpoint, it is not from a biological one. Each scenario answers to a specific question and changing the scenario implicitly amounts to changing the question one answers to. Second, it may suggest to unexperienced readers that it makes sense to test several statistical workflows and then to a posteriori choose the one which results are the most satisfactory. However, doing so is essentially equivalent to throwing the dice several times and choosing the best outcome. Such type of “hazard-cheating” is classically referred to as p-value hacking and it does not comply with scientific good practices. Nonetheless, when it comes to open the black box of statistical testing, relying on a concrete example to exhibit different behaviors is insightful. To cope with this dilemma, the chosen dataset corresponds to an experimental design which processing could advantageously rely on a two-way ANOVA, while the scenarios presented are centered on the one-way ANOVA. By doing so, one easily circumvents the question of which scenario to use “to best make the data speak” (in a way that would possibly differ from the scientific motivations of the authors who published the dataset in the first place) as to focus on the computational behavior of the statistical procedures.

The chosen dataset, hereafter referred to as RESET, results from a label-free proteomics experiment, which takes part into a larger multi-omic investigation about controlling the growth of *Escherichia coli* cells thanks to an external control of RNA polymerase expression [12]. It contains 6 different biological groups (3 samples each) and 1,634 identified and quantified proteins. The relationship between the six conditions is purposely ignored, and we will hereafter assume that all the pairwise comparisons are equally interesting. Doing so will lead us to consider the Tukey Honest Significant Differences (or Tukey HSD) as the running example of PHT. This PHT is probably the most straightforward to picture, and in addition, it makes the row-wise MTC particularly important as it leads to considering as many as 15 contrasts on 6 conditions. It has therefore a real pedagogical interest; however, replacing it by another PHT would not qualitatively alter the conclusions of the comparisons, nor the discussion of the article.

In the following sections, all the R codes to run each scenario on the RESET dataset are provided in the text. To have these code chunks run, it is necessary to rely on a recent version of R (version 4.3.0 or newer), as well as to have the following packages installed, together with their dependencies: **DAPAR, cp4p, readxl, multcomp, MSnbase, apcluster, Mfuzz, magrittr**. Then, it is necessary to download the data spreadsheet of [12] (available here: https://pubs.acs.org/doi/suppl/10.1021/acssynbio.1c00115/suppl_file/sb1c00115_si_006.xlsx) and to run the preliminary code provided in Supporting Information (see *suppmat1_R_code_RESET_data_preparation*.*rtf;* just copy-paste it in R console). This code opens a window to select the spreadsheet, imports its numerical data into the R environment and formats them as to make subsequent statistical processing the easiest possible.

To describe the scenarios, the following generic notations are used: *N* refers to the number of proteins being tested; *n* (respectively, *n*_*q*_) refers to the number of proteins being selected, globally (respectively, in contrast *q*); *Q* denotes the number of contrasts (which should not be confused with *K*, the number of groups or of biological conditions, see Figure 1a). To illustrate this on the RESET dataset, *N*=1634, *K*=6 and *Q*=15, while *n* and *n*_*q*_ (with *q* ≤ 15) will change depending on each scenario.

**Figure 1:**
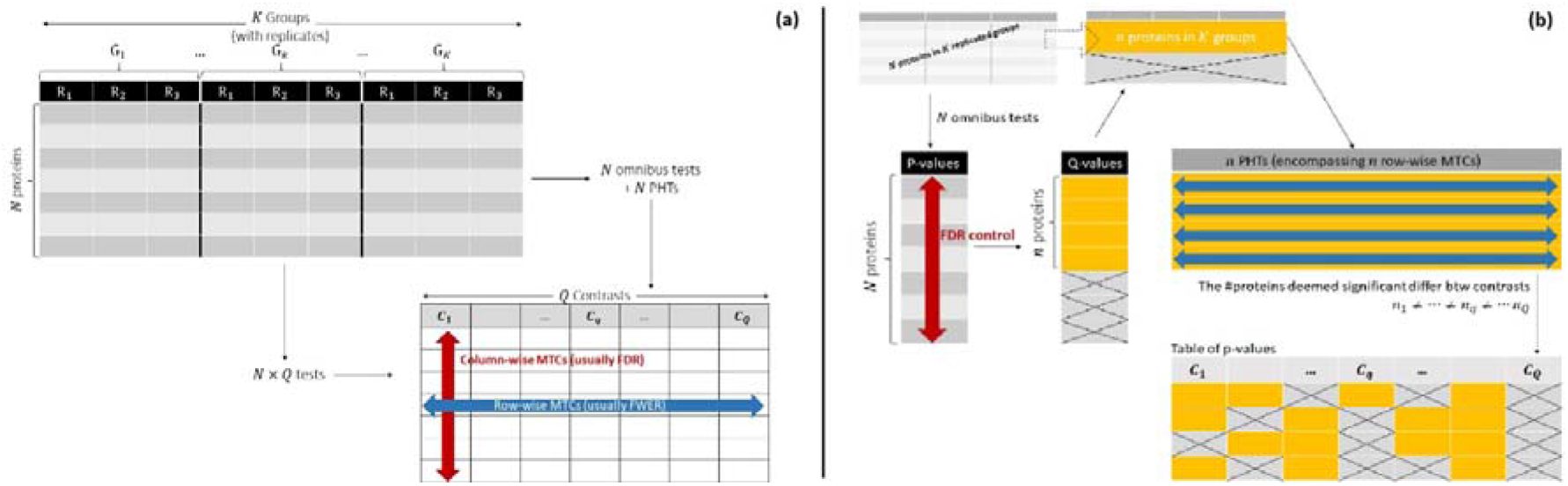
(a) schematic representations of the data and contrast tables with a recap of the mathematical notations. (b) The textbook scenario (omnibus tests + FDR + PHTs)

### 3.1. The textbook scenario: omnibus tests + FDR + PHTs

The most natural way to apply the ANOVA framework to the omics world is simply to iteratively process each omic features. In other words, if there are *N* quantified proteins, apply *N* omnibus tests, and for those which are deemed significant according to some criterion, apply a PHT. As in the proteomics community, the significance criterion for differential abundance analysis has long been the FDR (cf. Introduction), it also makes sense to use it to select the proteins worth of a PHT. Therefore, the complete procedure becomes: apply *N* omnibus tests on the *N* proteins, select the *n* ones with the smallest p-values, with *n* so that the FDR is controlled at a user-defined threshold, and then apply the PHTs on the original dataset (in fact, on its linear model), yet restricted to these *n* proteins only (see Figure 1b). Applying this scenario to the RESET dataset requires a few lines of R code using DAPAR package [36]:

~~~
# Compute ANOVA models
anova.models <- t(apply(qData,1, OWAnova, conditions=as.factor(sTab$Condition)))
names(anova.models) <- rownames(qData)
# Perform Omnibus test
omnibus.res <- testAnovaModels(anova.models, test = “Omnibus”)
# FDR control: Filter out proteins with an adjusted p-value < 1%
omnibus.adjp <- adjust.p(omnibus.res$P_Value$anova_1way_pval)$adjp
omnibus.da.prot <- which(omnibus.adjp$adjusted.p < 0.01)
cat(length(omnibus.da.prot), “proteins are DA out of “, nprot,
     ”(i.e., “, length(omnibus.da.prot)/nprot*100, “%)", sep="”)
# Post-Hoc Test
ressc.textbook <- testAnovaModels(anova.models[omnibus.da.prot], test = “TukeyHSD”)
# Important question shows up here: rejection threshold on FWER adjusted p-values?
textbook.seltab <- compute.selection.table(ressc.textbook$P_Value, 0.01) # why not 0.05?
textbook.seltab
~~~

The first chunk of code computes the linear model and the second one applies the omnibus test in a rather automated way. In the third chunk, one has to tune the FDR threshold. Rather classically, we can really on 1%, as from a QC perspective, it is the most standard threshold. However, in the next chunk, one has to tune the rejection threshold for the PHT, which will apply to the FWER-corrected p-values of the pairwise contrasts. Depending on the chosen value (as there is no “standard” one), the number of protein/contrast pairs exhibiting differential abundance will be different. As an example, at 1%, they are 9,541 of them, compared to 11,174 at 5% (+17%). Finally, the overall pipeline is extremely sensitive to the PHT significance level, so that the QC role classically attributed to the FDR becomes meaningless.

Moreover, it is important to understand that subsequently to the PHTs, some proteins may display differential abundance for zero, one or even several contrast(s). Conversely, the number of differentially abundant proteins may largely differ from one contrast to another. This can be easily pictured on the lower data table of Figure 2b, where the proportion of crossed cells within each column and in the entire data table is not the same. From the researcher viewpoint, this may compromise the PF role of the feature selection process, as it is not possible anymore to keep the desired number of candidates for post-proteomics validation associated to each contrast.

**Figure 2:**
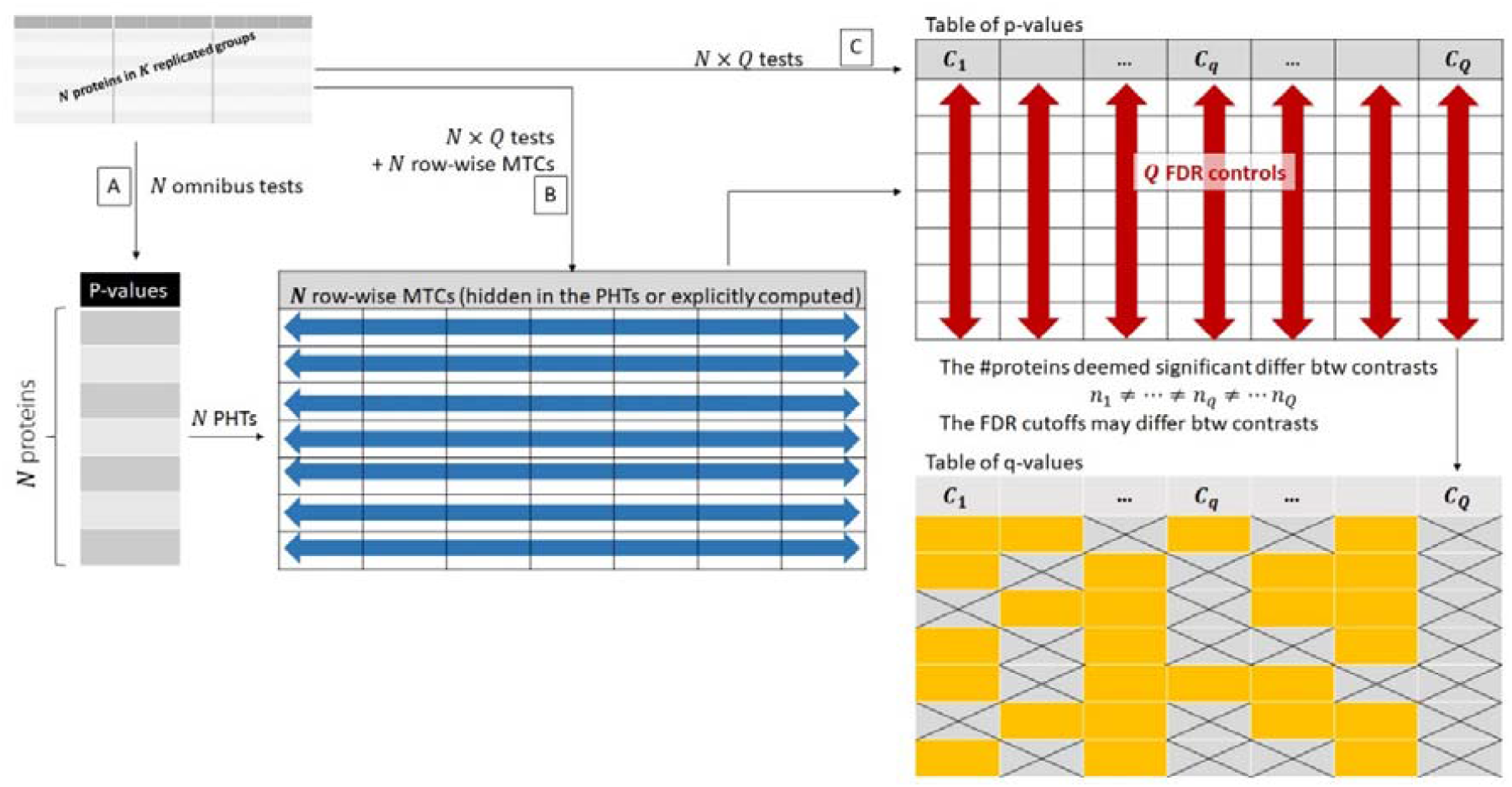
[A] The swap scenario (Omnibus tests + PHTs+ FDRs); [B] The contrast-based equivalent to the swap scenario; [C] The scenario corresponding to the “separate” option in Limma

Finally, the number of discovered differential abundances is strictly speaking not FDR-controlled, as each contrast was not corrected for the multiplicity of the protein tested (observe that on Figure 2b, after the FDR control, one goes back to a truncated version the original dataset, which has not undergone column-wise MTC). Therefore, if one protein exhibits a strong differential abundance in a first contrast (say group 1 vs. group 2), the low p-value of the omnibus test will make the protein pass the FDR threshold. Then, the protein can appear differentially abundant in other contrasts (say group 1 vs. group 3 or group 2 vs. group 3), where it should not have passed the significance threshold with an independent (column-wise) p-value adjustment. In other words, the SR role of the FDR is lost (in spite of a selection process, which can be tuned to be stringent enough).

To summarize, with this scenario, the FDR thresholding cannot fulfill with any of the three roles one expects. Considering, it should not be promoted for most of the application cases. However, as an easy-to-reproduce workflow, it can be interesting to reduce the number of hypotheses to subsequently explore (originally, as many as *n* times *Q*) when a proteomic experiment is conducted at a very exploratory stage of a research project. For instance, if one is primarily interested in a hand-waving overview of which proteins could be changers between various groups, so that an up/down regulation mechanisms can be hypothesized, upstream more comprehensive experiments and associated significance computations. However, as described below (see Section 3.3), another workflow is generally more adapted to do so.

### 3.2. Swapping scenarios: the FDR is saved for the end

An alternative to the previous scenario is to swap PHTs and FDR control. As described on the path A illustrated on Figure 2, the *N* omnibus tests are followed by as many PHTs, leading to a table of *Q* by *N* p-values, and *Q* FDR controls are finally applied to provide a subset of *Q* lists of respectively *n*_1_ *n*_2_, …, *n*_*Q*_ proteins. As with the previous scenario, the values *n*_1_ *n*_2_, …, *n*_*Q*_ may not be equal, and even in case of equality, the *Q* lists of differentially abundant proteins may not be the same from one tune the *Q* FDR thresholds to have broadly equal *n*_1_ *n*_2_, …, *n*_*Q*_ Although questioning from a SR contrast to another. However, and in contradiction with the previous scenario, the practitioner can viewpoint, doing so may be useful in a PF aim. Nevertheless, if the FDR thresholds are all tuned to a same value dictated by QC standards, then the scenario is compatible with both the SR and QC roles (briefly, as no processing occurs downstream of the FDR control, it remains valid).

This second scenario can be seen as the ultimate one, in the sense that it is sufficiently flexible to manage a trade-off between the three roles of FDR. However, studying the code to run it should raise questions:

~~~
# Compute ANOVA models
# Does not need to be recomputed after processing the textbook scenario
anova.models <- t(apply(qData,1, OWAnova, conditions=as.factor(sTab$Condition)))
names(anova.models) <- rownames(qData)
# Perform Omnibus test
# Does not need to be recomputed after processing the textbook scenario
omnibus.res <- testAnovaModels(anova.models, test = “Omnibus”)
# Post-Hoc Test
# NB: results from omnibus test is no longer used, previous step can be bypassed.
ressc.swap <- testAnovaModels(anova.models, test = “TukeyHSD”)
# FDR control: Filter out proteins with an adjusted p-value < 1%
swap.sep.padjtab <- separateAdjPval(ressc.swap$P_Value)
swap.sep.seltab <- compute.selection.table(swap.sep.padjtab, 0.01)
swap.sep.seltab
~~~

The first two code chunks are the same as in the previous scenario, so they do not need to be re-run. As for the third one, interestingly enough, it does not take the omnibus test result as input (second chunk), but the linear model (first chunk). It means the omnibus test chunk is in fact useless for this scenario. As argued in Sections 2.3 and 2.4, if an omnibus test is systematically followed by PHTs, the omnibus test is not needed (at least, no longer, as saving computations could be a valuable objectives few decades ago). Conversely, a workflow like “Linear model + multiple contrast tests + protein-wise FWER controls + contrast-wise FDR controls” (as illustrated on the path B of Figure 2) is more representative of the R code used.

As this workflow adheres to Limma’s philosophy (see Section 2.3), let us check if it is possible to run it while benefiting from the moderated statistics. In the documentation, four options are implemented (see the `decideTests` function, [35]). Those termed `“separate”` and `“global”` do not correct for contrast multiplicity and are discussed in the next section. The `“NestedF”` one is restricted to microarray data. Finally, the `“hierarchical”` one is the closest from our expectations, in the sense that it proposes both row-wise and column-wise MTCs. Unfortunately, the documentation contains the following warning: “The `“hierarchical”` *method offers power advantages when used with* *adjust*.*method=“holm”* to control the family-wise error rate. However its properties are not yet well understood with `adjust=“BH”`.” In other words, the MTC can either be an FWER with statistical guarantees (Holm’s method amounts to a sequential Bonferroni-like control of the FWER [27]), or an FDR, without statistical guarantees (BH refers to Benjamini and Hochberg procedure to control the FDR [18]). Moreover, a deeper look in the code indicates the chosen MTC (be it Holm or BH) is applied both row-wise and column-wise, so that it does not fit our scope, where the column-wise MTC is expected to be an FDR and the row-wise one an FWER.

### 3.3. The no-FWER scenarios

Boldly skipping the row-wise MTCs in the previous scenario (as illustrated on the path C of Figure 2) may be tempting, as doing so will avoid the inflation of p-values that is subsequent to the FWER control:

~~~
# re-test the same anova models without MTC
result.noFWER <- testAnovaModels(anova.models, test = “TukeyNoMTC”)
# directly apply FDR control (1%)
noFWER.sep.padjtab <- separateAdjPval(result.noFWER$P_Value)
noFWER.sep.seltab <- compute.selection.table(noFWER.sep.padjtab, 0.01)
noFWER.sep.seltab
~~~

Indeed, more protein/contrast pairs appear as differentially abundant using this approach than the previous one involving an FWER control (11,030 vs. 8,610). At first glance, this gain of statistical power may appear as a p-value hacking, which is incompatible with statistical rigor. However, it is not necessarily the case if one avoids subsequent over-interpretation. In fact, this scenario is rather common: it is implemented in Limma (under the method `“separate”` in the `decideTests` function) where it is the default method. As explained in the documentation, this method “[…] *does multiple testing for each contrast separately […]. Using this method, testing a set of contrasts together will give the same results as when each contrast is tested on its own. The great advantage of this method is that it gives the same results regardless of which set of contrasts are tested together. The disadvantage of this method is that it does not do any multiple testing adjustment between contrasts. Another disadvantage is that the raw p-value cutoff corresponding to significance can be very different for different contrasts […]. This method is recommended when different contrasts are being analysed to answer more or less independent questions*.” Using DAPAR routines, the Limma version of this scenario reads:

~~~
# and Limma counterpart
limma.res <- limmaCompleteTest(qData, sTab, comp.type="OnevsOne”)
# subsequent FDR control
limma.sep.padjtab <- separateAdjPval(limma.res$P_Value)
limma.sep.seltab <- compute.selection.table(limma.sep.padjtab, 0.01)
limma.sep.seltab
~~~

Using this code, the number of differential protein/contrast pairs is even larger (11,281). The increment with respect to the previous one (11,030) directly stems from Limma’s moderated statistics (see Discussion). Anyway, using this scenario, the most difficult point is to determine whether the questions at stake are sufficiently independent to justify the suppression on the FWER control. Notably, comparisons involving independent biological conditions do not imply that the associated contrasts (or questions) are independent. As a counter-example, just think about the testing of hundreds of placebo treatments depicted in the introduction: although chemically independent, the manifold of treatments requires MTC. Finally, this scenario trades some statistical prudence with a small increment of the statistical power: the only difficulty is to determine whether this imprudence is acceptable considering the data, or if it hides some data dredging. In the second case only, the SR role attributed to the FDR is compromised by the lack of upstream FWER control. As for the PF and QC roles, this scenario stands exactly at the same point as that of Section 3.2.

Let us now depict another situation where it is possible to solve the entanglement of row-wise FWERs and column-wise FDRs by getting rid of the former ones, without compromising the SR role of the latter ones. The corresponding scenario is well suited to answer a “global” question, *i*.*e*., a question that does not translate into several specific contrasts. In those cases, the statistical rigor is preserved despite the absence of FWER control because multiple contrasts are no longer tested simultaneously. As example, consider a time-course with so many time-stamps that exhaustive pairwise comparisons are intractable. To cope for this, one focuses on the global trend of each protein, as to discriminate between those which abundance remains stable despite random fluctuations, and those which abundance changes significantly. Of course, one may also be interested in whether each variation of abundance amounts to an up or down-regulation; as well as whether it occurs at the beginning, at the middle or at the end of the time-course. However, depending on the application, these refinements may remain of qualitative nature (*i*.*e*., devoid of associated significance values), without flawing the biological reasoning and results.

In those cases, the omnibus test is sufficient to provide a global significance value to each protein profile, while complementary information can be emphasized with exploratory and descriptive tools, like cluster analysis. As an illustration, Figure3a schematizes several clusters obtained on averaged and standardized protein profiles (*i*.*e*., each protein profile is constructed by averaging all the samples’ values within each group; then the profile is centered on 0 and scaled to a unitary variance [37]). By combining these clusters with the result of an omnibus test, it is possible to spot proteins which profiles (whatever their shape) are significantly different from a flat profile, according to the user-defined FDR threshold. Doing so using DAPAR routines is rather straightforward:

~~~
omnibus.limma.res <- limmaCompleteTest(qData, sTab, comp.type="anova1way”)
wrapperRunClustering(obj = MSnSet(exprs = as.matrix(qData), fData = qData, pData = sTab),
                 adjusted_pvals =
                   adjust.p(omnibus.limma.res$P_Value$anova_1way_pval)$adjp$adjusted.p,
                 clustering_method = “kmeans",
                 k_clusters = 9,
                 FDR_thresholds = c(0.01))
~~~

The code can be customized in many ways to change the number of clusters, the clustering method, the omnibus test (e.g., replacing Limma’s version by the classical ANOVA one) or the FDR cutoffs used to highlight protein profiles with significant trends. For most proteomic experiments that will need post-analytics validations, results of this type are sufficient. However, it is important to avoid over-interpretation when describing them. For instance, if among a cluster exhibiting an early downfall in profiles, one protein is significantly differentially abundant according to the omnibus test, it is not possible to conclude that the early down-regulation itself is significant. Saying so would amount to transfer the significance associated to the global rejection of 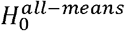 to a more specific question (namely, the early down-regulation), which would constitute a statistically unsupported claim. However, it remains nonetheless possible to conduct downstream experiments to assess further this early down-regulation. Therefore, as long as one discriminates between conclusions that require statistical significance supports, and intermediate evidence along the course of a biological investigation (which may not require a similar support as experimental confirmations are expected), it is possible to keep a safe and rigorous line of reasoning. Most importantly, doing so makes it possible to reconcile the three roles of FDR control: SR (as long as one does not over-interpret and accepts an answer to a broader question), PF and QC (notably, managing a single selection process makes it easier to reach an agreement between the last two roles, as with a single binary contrast).

Lastly, in the cases which do not fall in the previous categories (*i*.*e*., the questions are not sufficiently independent and they require going beyond a global approach), the simplest workaround is to stack all the contrasts in a single large contrast, so that the cumbersome combinations of row-wise and column-wise MTCs is replaced by a single column-wise MTC (see Figure 3b). Concretely, two p-values yielded by a same protein for two different contrasts are considered exactly as two p-values resulting from two proteins tested in a single contrast. This strategy has also been promoted by Limma’s authors, through the method referred to as “`global`” in the `decideTests` function. This approach has many advantages, which are listed in Limma user’s guide [35]. Their paraphrasing here would be of little interest, but to summarize, the PF and QC roles classically endowed to FDR control procedure are fully accounted for.

**Figure 3:**
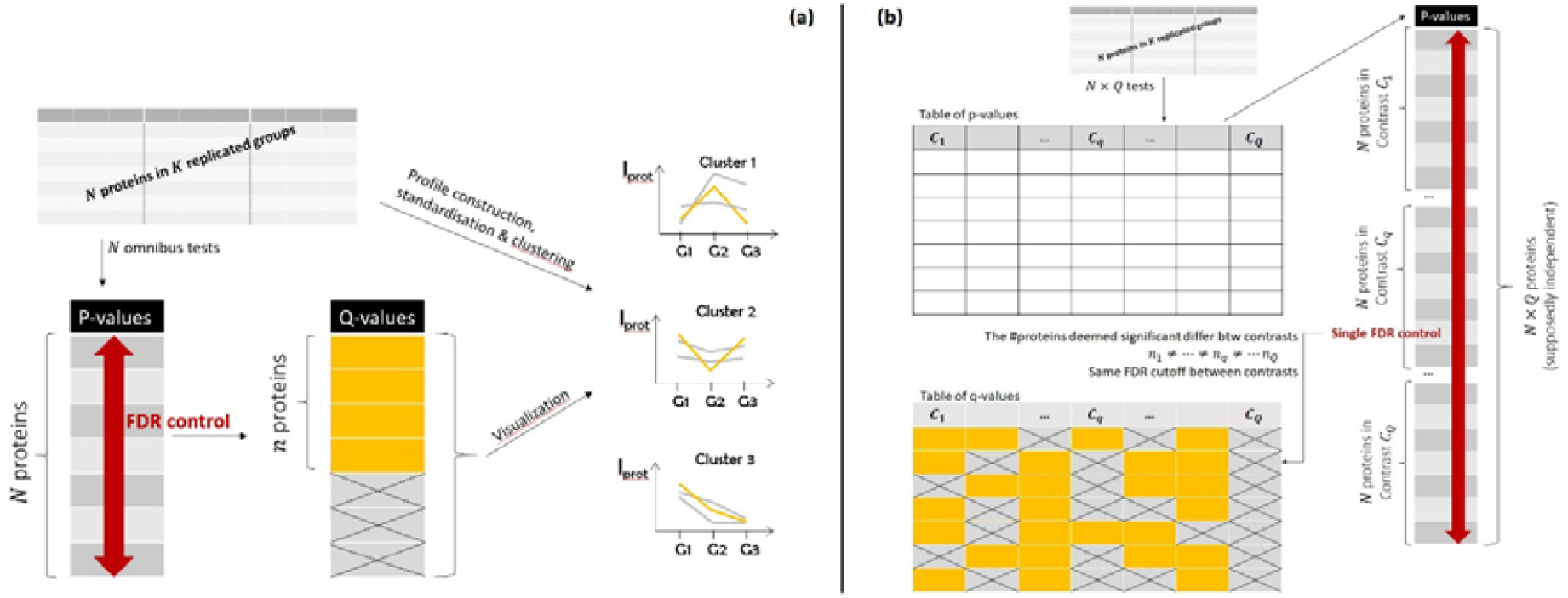
(a) Omnibus tests + FDR + Clustering; (b) Single FDR control over stacked contrasts (“global” option Limma)

Unfortunately, depending on the column-wise MTC, this approach may lack theoretical support. Notably, when this MTC amounts to an FDR control (which is the most classical approach in proteomics), the authors say the following: “[…] *there is no theorem which proves that* *adjust*.*method=“BH”* in combination with `method=“global”` *will correctly control the false discovery rate for combinations of negatively correlated contrasts, however simulations, experience and some theory suggest that the method is safe in practice*.” Although the absence of theoretical guarantee should not systematically discard the application of an otherwise interesting approach, relying on qualitative and simulation-based empirical arguments only is not sufficient to trust in the SR role fulfillment. A middle of the road approach is to stack the *Q* p-value vectors manually and then to run a tool like CP4P [38]. CP4P makes it possible to check the conditions of application of BH FDR control through the visualization of the p-value calibration, as well as to some extent, to correct for minor calibration defaults, as detailed in [24]. Then, depending on the calibration assessment, the practitioner may decide (or not) to use this scenario (*i*.*e*., a single FDR control on stacked contrasts, *i*.*e*., `adjust.method=“BH”` in combination with `method=“global”` in Limma). The following code implements this strategy:

~~~
# Same linear model previously computed
limma.res <- limmaCompleteTest(qData, sTab, comp.type="OnevsOne”)
# Calibration check: In this case, it is nearly perfect!
calibration.plot(stack(limma.res$P_Value)[,1])
# FDR control
limma.glob.padjtab <- globalAdjPval(limma.res$P_Value)
limma.glob.seltab <- compute.selection.table(limma.glob.padjtab, 0.01)
limma.glob.seltab
~~~

The first code chunks re-compute the linear model as used before. The second chunks display the CP4P plot, which in this case, indicate a very good calibration. Then, the last one deals with the FDR, like in the previous scenarios.

### 3.4. More advanced scenarios

The above scenarios should answer most experimental designs. However, it is possible to tailor more specific tools based on another generic approach: Firstly, for each protein, one aggregates all the p-values resulting from the *Q* contrasts into a single p-value; secondly, one applies an FDR control on this single “aggregated contrast” (see Figure 4). Theoretically speaking, aggregating the multiple p-values resulting from multiple contrasts into a single protein-wise p-value can be seen as a form of MTC. Thus, by leveraging the appropriate procedure, the SR role should be preserved.

**Figure 4:**
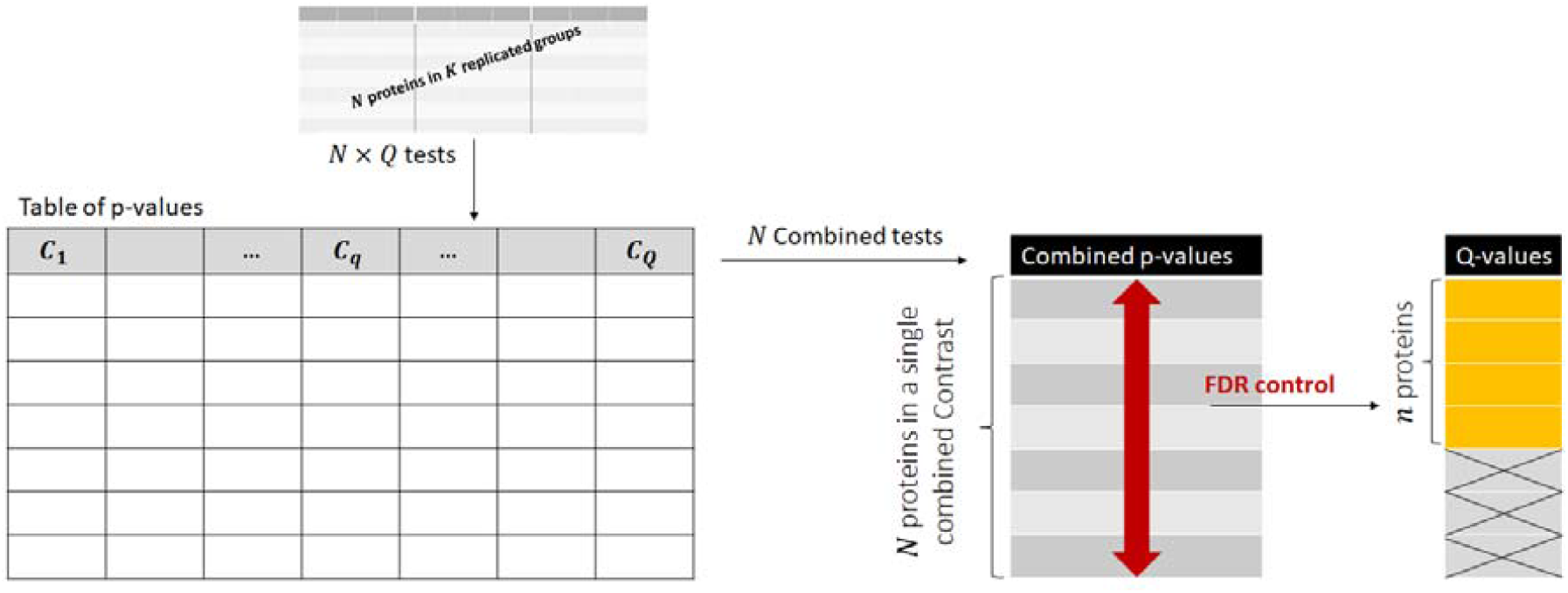
In this scenario, the various contrast p-values are aggregated for each protein so as to provide a single p-value for each, then FDR control is applied.

Many p-value combination methods endowed with proof of correct calibration exist in the literature (Harmonic mean [39], Fisher combination [40], etc.), but not as many can be interpreted as contrast summarization. Following a previously described use-case, let us assume we are mainly interested by a global view of differential abundance, as for instance, in a time-course experiment. In this context, a protein can be interesting because it displays a consistent trend, even if this trend is sometimes significant and sometimes not (for instance, the protein is constantly up-regulated, but this is not significant between some consecutive time stamps, while it is between others). One may thus be tempted to focus on the smallest p-value between the *Q* contrasts for each protein, as a marker of its “most significant fold change”. However, taking the minimum out of *Q* p-values requires a subsequent Šidák correction [41], which as-a-matter-of-factly corresponds to an FWER control:

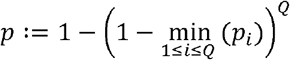

After computing this single p-value for each protein, it becomes easy to subsequently control the FDR, as a single “aggregated contrast” remains. Conversely, if one is interested in the “worst fold change” among *Q* contrasts, it is possible to consider the max p-value. Although very conservative, this can be instrumental when comparing the weakest link among cascading events. Then, it is possible to adapt Šidák correction to maintain the correct calibration, and to compute p-values as follows:

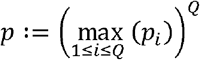

The interest of this approach is that depending on the question at stake, it is possible to rely on the important body of literature about how to combine statistical tests to simplify the interferences between the MTCs resulting from the multiple contrasts and those resulting from the manifold of proteins tested. For illustration purpose, the R codes to apply these worst / best-case scenarios are provided in Supporting Information (either with a classical ANOVA model, or with Limma, see file *suppmat2_R_complete_code_for_exhaustive_comparisons*.*rtf*).

Likewise, it is possible for advanced users to mix the scenarios of Sections 3.1 to 3.3. For instance, it is possible to apply the “global” method (stack all the contrasts prior to FDR control) inherited from Limma to a classical ANOVA. The combinatory of all these “mixed scenarios” is rather high, so the code chunks are provided in the Supporting Information (file *suppmat2_R_complete_code_for_exhaustive_comparisons*.*rtf*). Using this code, it is possible to compare the amount of proteins deemed differentially expressed according to each scenario. Doing so to choose the one providing the largest amount of proteins is of course nonsensical from a statistical viewpoint. However, a global comparison of their result sheds an interesting light about the conservative/liberal behavior of each component of the workflows, as detailed in the next section.

## 4. Discussion

Using the codes available in Supporting Information, Table 1 summarizes various scenarios with respect to the number of significant (protein, contrast) pairs. This comparison leaves aside the textbook scenario (Section 3.1), but to define a baseline, let us recall that if its PHT significance threshold is moved from 1% to 5% (irrespective of the FDR threshold), 17% of additional significant contrasts are found (from 9,541 to 11,174). With this in mind, let us compare the “global” and “separate” approach to FDR, the moderated and classical linear models (i.e., using Limma or a classical ANOVA), and 2 scenarios (the “swapped” one —see Section 3.2; and the No-FWER one —see Section 3.3). However, let us recall that Limma does not propose a scenario like the swapped one, and forcing its implementation would be hazardous, because of the lack of theoretical and empirical studies about the interference between two different types of MTCs and the moderated statistics.

**Table 1:** Number of significant (protein, contrast) pairs according to the various scenarios

|  | SEPARATE | GLOBAL |
| --- | --- | --- |
| **SWAP (CLASSIC)** | 8,610 | 8,586 |
| **NO-FWER (CLASSIC)** | 11,030 | 11,077 |
| **NO-FWER (LIMMA)** | 11,281 | 11,349 |

Based on the numbers in the columns, it appears that using a global or separate FDR control has little influence on the conservative/liberal behavior of the workflow. As advocated in Limma’s documentation, it is essentially a matter experiment design (independence of the contrasts), of choice (each approach having their pros and cons, described in the documentation) and of correct calibration. However, comparing the rows of the table is more insightful. Broadly, using the moderated statistics, 250 to 275 additional contrasts are found significant (i.e. the difference between the last two lines). Conversely, adding an FWER control leads to 10 times more contrasts (≈ 2,500) being considered as non-significant. This clearly rise questions about Limma’s increased sensitivity, which is the main reason of its popularity. While it partially roots in the moderated statistics (as regularly advocated), one sees it also roots in the lack of MTC across contrasts. In fact, this latter reason is majority by a tenfold. A similar comparison about the best/worst contrast scenarios (Section 3.4) is reported in Table 2, and it also indicates that the moderated statistics, although interesting, is not a game-changer on its own. On the contrary, the absence of row-wise MTC (or a change in the PHT threshold in the case of the textbook scenario) can dramatically increase the number of proteins that are declared differentially abundant. However, both must be scrutinized at the light of possible p-value hacking. With this regards, it has been recently demonstrated that excessive trust in moderated statistics could be damaging [7], and guidance about when and how row-wise MTC can be safely gotten rid of (and when it cannot) as provided here, is insightful.

**Table 2:** Number and percentage of proteins deemed significant (out of 1,634) according to their best and worst contrast with Limma and a classical linear model

|  | WORST CONTRAST | BEST CONTRAST |
| --- | --- | --- |
| **LIMMA** | 353 (21.7%) | 1,334 (81.6%) |
| **CLASSIC MODEL** | 354 (21.7%) | 1,303 (79.7%) |

## 5. Conclusions

The fixed-effect one-way ANOVA workflow is particularly popular to process omics data. The reasons are threefold: First, its genericity, which makes it suitable enough to a large variety of designs and of biological questions. Second, its ease of understanding and interpretation, as a straightforward extension to the t-test. Third, its considerable importance from a historical perspective.

However, its systematic use may not be adapted to modern omics data: First, despite being rather generic, it is usually possible to find an alternative contrast-based workflow that is more specific to the design or biological question considered. Second, its computational efficacy has become of limited (or null) interest according to the modern computational resources. Finally, its PHTs difficultly orchestrate with the need for FDR control. All these advocate for a restricted use of the traditional one-way ANOVA workflow, and for the growing use of now well-established contrast-based scenarios.

Doing so notably makes it possible to replace ANOVA PHTs by off-the-shelf FWER control procedures. The latter ones are generally a bit simpler to combine with FDR controls, even though complex situations may still exist. For those, it is mandatory to wonder about which roles are expected from the FDR control, between “statistical rigor”, “practical filter” and “quality control”: Depending on their respective importance, one or several scenarios among those listed above can be preferred.

## Supporting information

suppmat1_R_code_RESET_data_preparation.rtf

suppmat2_R_complete_code_for_exhaustive_comparisons.rtf

## Acknowledgement and fundings

The author wish to thank Hélène Borges, Yohann Couté and Virginie Brun for fruitful discussions. This work was supported by grants from the French National Research Agency: ProFI project (ANR-10-INBS-08), GRAL project (ANR-10-LABX-49-01), LIFE project (ANR-15-IDEX-02) and MIAI @ Grenoble Alpes (ANR-19-P3IA-0003).

## Footnotes

1 Applying a cut-off on the fold-change is a common practice in proteomics, however, if improperly achieved, it can hinder the statistical rigor. Notably, it has been demonstrated that applying such a cut-off downstream of the FDR is invalid [4]. It remains possible to incorporate the fold-change cut-off in the statistical test [22], but the resulting p-value interpretation can be a bit more difficult. Moreover, depending on how it is conducted, it can even lead to involuntary p-value hacking [23], so that applying such a filter prior to differential abundance testing constitutes a nice trade-off [24].

